# Mechanisms underlying higher order interactions: from quantitative definitions to ecological processes

**DOI:** 10.1101/857920

**Authors:** Andrew R. Kleinhesselink, Nathan J.B. Kraft, Jonathan M. Levine

## Abstract

When species simultaneously compete with two or more species of competitor, higher order interactions (HOIs) can lead to emergent properties not present when species interact in isolated pairs. In order to extend ecological theory to multi-competitor communities, ecologists must develop a practical and general definition for HOIs that can be applied to a wide range of competition models. In this paper we propose a definition for HOIs and outline a set of criteria for testing whether a model has or does not have HOIs. These criteria are valuable for empirical ecologists in need of clarity when discussing HOIs in empirical data. We also provide thorough discussion of how our definition compares with previous definitions of HOIs and interaction modification in the literature. In the second part of the paper we demonstrate the steps required for a rigorous test of HOIs in empirical data. To do this we simulate resource competition between three annual plant species which differ in phenology. We then fit phenomenological competition models to the outcome of simulated competition and use these to test for the presence of HOIs. In our simulations, we find the strength of HOIs varies with phenology: species that grow later experience stronger HOIs than earlier growing species. Our simulation shows how HOIs could emerge in ecosystems where resource availability and individual size change rapidly throughout the course of the growing season and where there are differences in the timing of resource acquisition between competitors.

## Introduction

Almost all species interact with a diversity of predators, pathogens and competitors. Despite this, most classical models in community ecology assume that the per capita effects of each species on each other do not dependent on the densities of any other species in the community. This simplifying assumption means that we can predict the dynamics of multispecies communities from a model that only includes the interaction between each pairs of species (Chesson 2000, Levine et al. 2017).

Higher order interactions (HOIs) between species invalidate the core assumption of independent per capita interactions and thus HOIs could have profound consequences for modeling community dynamics and species coexistence (Neill 1974, Mayfield and Stouffer 2017, Levine et al. 2017, Grilli et al. 2017). If HOIs are strong, even a perfect understanding of the interaction between each and every pair of species in isolation would not be sufficient to describe what happens when all the species are simultaneously interacting (Neill 1974, Billick and Case 1994, Levine et al. 2017). A specific example of the potential for HOIs to impact our understanding of community dynamics is in the application of the mutual invasibility criterion for determining the stability of coexistence (Levine et al. 2017). In theory, HOIs can allow three competitor species to coexist even where some pairs of competitors cannot coexist (Grilli et al. 2017).

Despite the theoretical importance of HOIs, measuring HOIs in nature has been impeded by shifting definitions of what does and does not count as an HOIs (Pomerantz 1981, Adler and Morris 1994, Billick and Case 1994, Letten and Stouffer 2019). Moreover, previous definitions of HOIs were developed with a small range of classical competition models in mind. Since that time, new statistical modeling software now allows ecologists to fit a much wider range of interaction models (Mayfield and Stouffer 2017). This increase in model flexibility requires deriving a more general definition for HOIs that can be applied to any density dependent model of population dynamics.

In addition, to the basic issue of producing a shared definition for HOIs, ecologists lack a mechanistic understanding of how HOIs could emerge in nature (Levine et al. 2017, Letten and Stouffer 2019). Such an understanding is necessary for predicting the sets of competitors and ecosystems where strong HOIs are likely. One promising way to address these outstanding issues is to simulate virtual competition experiments based on mechanistic models in which the processes that cause competition are fully known, and then evaluate for which species, and under which conditions HOIs emerge (Letten and Stouffer 2018).

We provide a general definition for HOIs based on interaction modification that distinguishes HOIs from related phenomena such as non-linear density dependence and indirect effects. In the second part of the paper, we use a simulation experiment to illustrate how our definition can be applied to properly identify interaction modification even against a backdrop of nonlinear density dependence. We then use the results of the simulation to shed light on possible mechanisms that could generate HOIs in nature.

### Higher order interactions result from interaction modification

For the purpose of defining HOIs we focus on modeling a focal species’ performance (usually per capita population growth rate) as a function of the population density of multiple species of competitor. This can be expressed generally as,

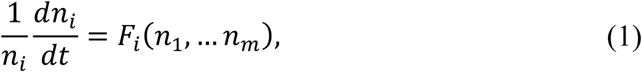

where *F*_*i*_ gives the per capita population growth rate of the focal species *i*, and *n*_*j*_ are the population densities of competitor species one through *m* in the community, including the population density of the focal species, *n*_*i*_. An analogous equation holds for population growth rate over discrete time intervals: 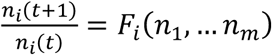. In most widely used models of species interactions, each competitor has one effect on itself and one effect on each of the other species in the community. The simplest example of such a pairwise competition model is the Lotka-Volterra (LV) model,

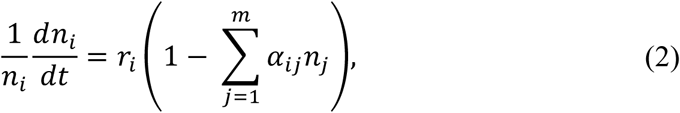

where, *r*_*i*_ is the intrinsic rate of growth of the focal species *i* and *α*_*ij*_ is the per capita effect of competitor *j* on the growth rate of the focal species. This model is pairwise because each interaction is specified by the pair of species involved, the focal species *i* and the competitor *j*. The defining property of any pairwise model, such as the LV model, is that the per capita effect of each species of competitor is independent of the densities of any *other* species of competitor (Figure 1A).

**Figure 1.**
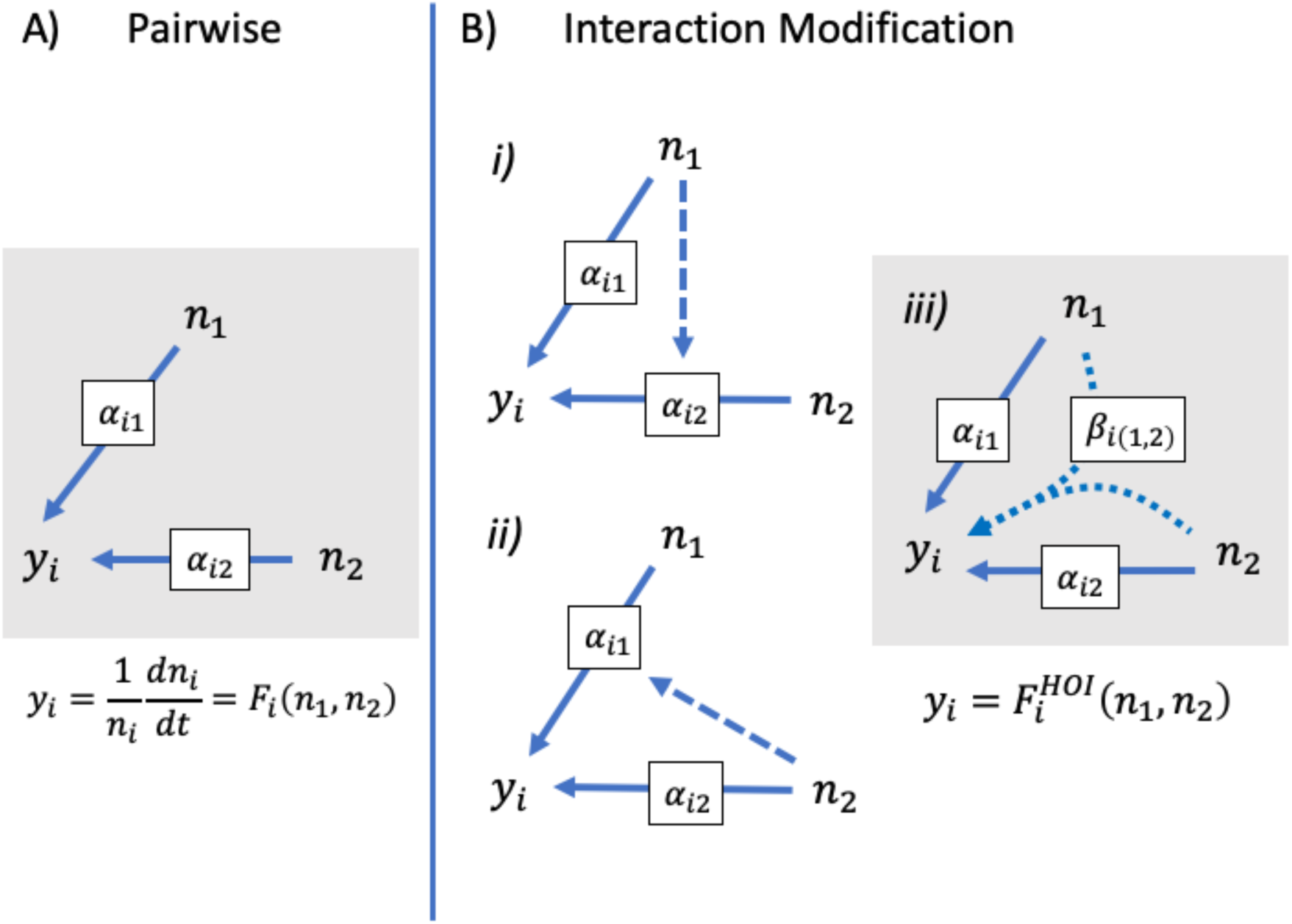
How interaction modifications lead to higher order interactions. In A, a pairwise model is shown without interaction modification. The competitive effect of species one and two on the per capita growth of the focal species *i*, are shown as separate blue arrows. These effects may be simple per competition coefficients, *α*_12_ and *α*_13_, or could be more complicated non-linear functions of density. In B, a model with interaction modification is shown: in *i)* the dashed arrow shows that the effect of two is modified by the density of one; in *ii)* the effect of one is modified by the density of two. In reality, one cannot assign either species as the modifier, rather they modify each other’s effects in a way that emerges a single HOI. The HOI in this case is quantified by introducing the new parameter *β*_*i*(1,2)_, and shown with the curved arrows in *iii*.

By contrast, interaction modification disrupts pairwise competition and leads to HOIs. Interaction modification occurs when the effect of one competitor species is modified by the density of another competitor species (Adler and Morris 1994). We can introduce an interaction modification into the LV model by replacing any of the constant terms *α*_*ij*_ with a function of the density of another competitor (Billick and Case 1994).

For instance, in the following LV model, the focal species performance is dependent on two competitor species,

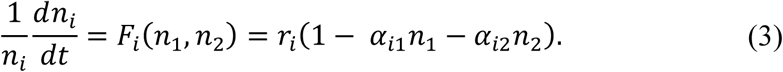

Replacing the term *α*_*i*1_ with the expression *α*_*i*1_ + *β*_*i*(12)_*n*_2_, makes the per capita effect of species one dependent on the density of another competitor, *n*_2_. More specifically the parameter *β*_*i*(12)_ measures the strength of this interaction modification (Figure 1B). Substituting this function into the model introduces the product of competitors one and two as a new term,

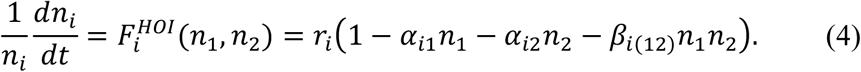

Interaction modifications such as these imply that competition is functionally different when more than one competitor species is present and that there are emergent properties in the community that cannot be predicted by single species effects. These may suggest specific biological hypotheses: something about the behavior or traits of the competitors are functionally disctinct when they are together as compared to when they are separate. Importantly, an interaction modification cannot be attributed to any one competitor— rather it is an emergent property of the multi-species system, what we call an HOI (Figure 1B).

### An improved general definition of HOIs

While the section above captures the essential connection between interaction modifications and HOIs, ecologists do not have a shared definition for HOIs that captures this idea and which can be applied to any density dependent model of competition (Hairston et al. 1968, Pomerantz 1981, Billick and Case 1994, Grilli et al. 2017, Letten and Stouffer 2019). Here we provide a formal mathematical definition for HOIs rooted in their important implications for ecological theory and which can be applied to any interaction model of any functional form. We first present this more formal definition but follow up with a simple empirical heuristic which can be used to evaluate a model for HOIs.

Let *F*_*i*_(*n*_1_, … *n*_*m*_) be a generic model describing the density dependent effects of *m* competitor species on the per capita growth of species *i*, where *m* > 1. Let Θ be the set of all parameters in the model, Θ = {*θ* | *F*_*i*_(*n*_1_… *n*_*m*_|*θ*)}. Here, the term parameter refers to constants in a model that are not themselves dependent variables (Bard 1974). Let *f*_*ij*_ (*n*_*j*_) be a model describing the response of the focal species to competition from a single competitor species, *j*, where *j* is one of the competitor species included in *F*_*i*_(*n*_1_, … *n*_*m*_). For any model *F*_*i*_, we find *f*_*ij*_ (*n*_*j*_) by setting the densities of all competitors except *j* to zero and simplifying the model. Next, let Ψ_*j*_ be the set of parameters in *f*_*ij*_ (*n*_*j*_), Ψ_j_ = {*ψ* | *f*_*ij*_(*n*_*j*_|*ψ*)}. For most realistic competition models the parameters in Ψ_j_ will be a subset of those in Θ, i.e. Ψ_j_ ⊆ Θ. Next, let Φ be the set of all parameters found across all *m* sets 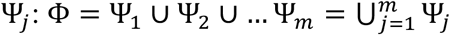. A model is pairwise if all parameters in Θ are found in the set Φ, i.e. Θ = Φ. **Models with HOIs are defined by having parameters in *F***_***i***_ **that are not found in the *m* single-competitor functions, or more precisely, when Θ is a proper superset of Φ, Θ** ⊃ **Φ**. Finally, let B be the set of parameters in Θ but not in Φ, B = Θ − Φ. The parameters in B are those that define the HOI in the model.

As a concrete illustration of our definition, consider the two competitor LV model defined in equation (3): for the full model Θ = {*α*_*i*1_, *α*_*i*2_, *r*_*i*_} and 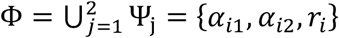, thus Θ = Φ and the model is pairwise. By contrast, for the HOI model 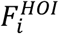 defined in equation (4), Θ = {*α*_*i*1_, *α*_*i*2_, *β*_*i*(12)_, *r*_*i*_} and 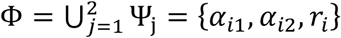, thus Θ ⊃ Φ and the model contains HOIs. Moreover, B = Θ − Φ = {*β*_*i*(12)_}, thus the parameter *β*_*i*(12)_ is specifically the one that captures the HOI.

This abstract representation belies a simple empirical heuristic for determining whether a model has HOIs: in order to parameterize a model with HOIs, the response of the focal species must be measured against density gradients of each competitor separately, as well as against varying combinations of competitors grown together (Figure 2). This is a natural consequence of the above definition. *In essence, a model with HOIs includes additional parameters that an empiricist cannot measure when the response of a focal individual is measured against a single competitor species (Pomerantz 1981)*. Note, however, there is no way to determine whether there are HOIs among *m* competitors by examining all *m* pairwise models *f*_*ij*_, rather the form for the multi-competitor model *F*_*i*_ must be chosen first in order to apply any HOI definition (Adler and Morris 1994).

**Figure 2.**
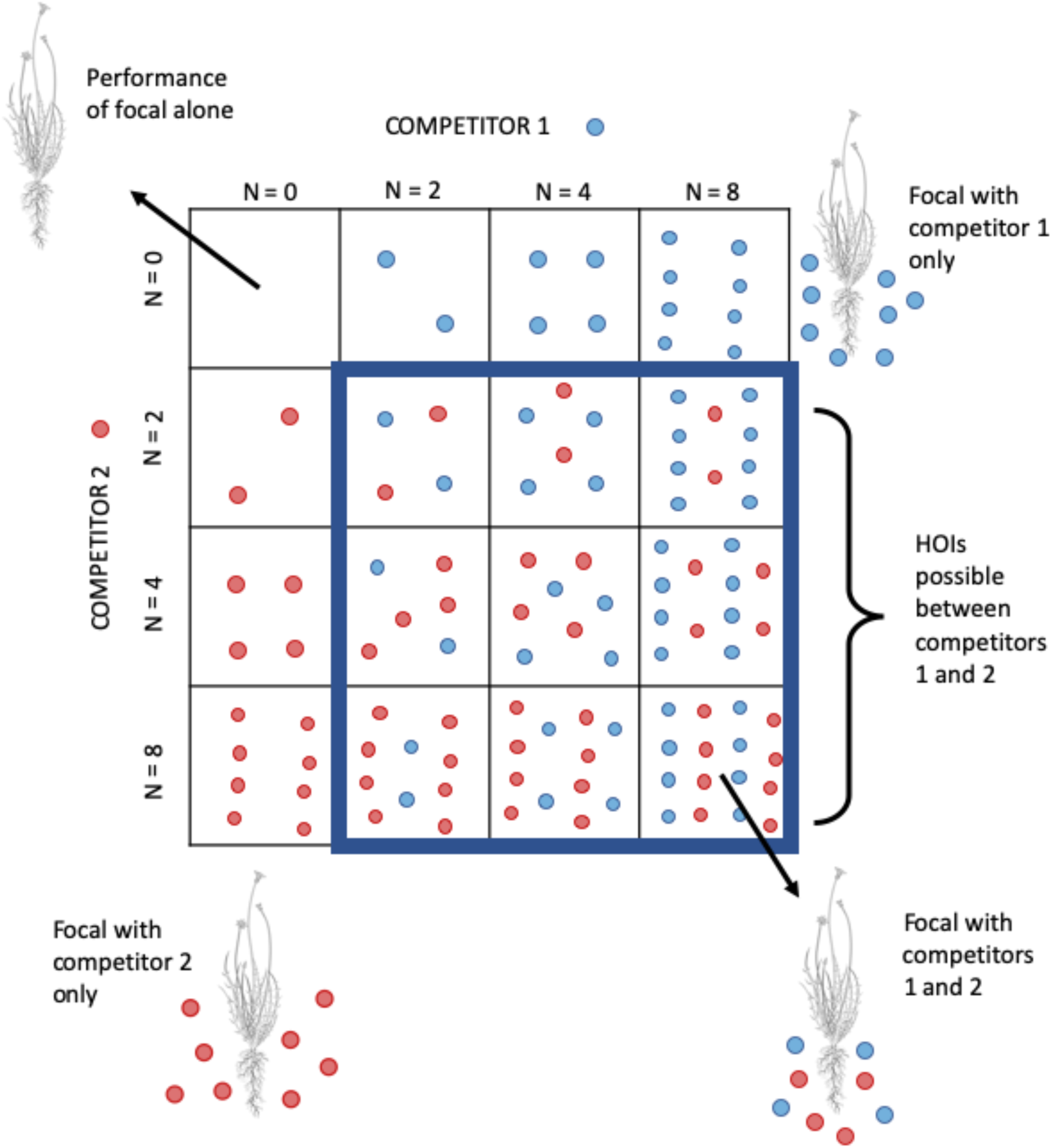
Schematic of orthogonal competition experiment required to detect higher order interactions. Each square represents a separate study plot. Competitor 1, (blue circles) and Competitor 2 (red circles) are grown in a gradient of increasing density alone and together. A single individual of the focal species (line drawing) is grown in each plot allowing the response to competition from each competitor species to be fitted.

We refer to the type of HOIs captured by our definition above as *hard HOIs* and contrast them with the wider phenomenon of non-linear density dependence which produces what we term *soft HOI*s. A general test for soft HOIs is to take the partial derivative of the competition function, *F*_*i*_ in equation (1), with respect to the density of a single competitor species, 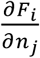. This partial derivative defines the focal species’ sensitivity to a single competitor. If this partial derivative is a function of more than one competitors’ density, then there are soft HOIs. In general, all models with hard HOIs will be non-linear and have soft HOIs, but not all non-linear models will have hard HOIs. This is similar to definitions used in earlier discussions of HOIs based on LV forms of competition (Case and Bender 1981), and closely follows the verbal argument that HOIs emerge when the effect of one competitor on another depends on any other competitors. The problem is that **any model in which growth is a nonlinear function of interspecific density will involve soft HOIs**, and thus this definition does not distinguish interaction modification or HOIs from non-linear density dependence (Pomerantz 1981, Adler and Morris 1994). As an example consider the multi-competitor Hassel model (Hassell and Comins 1976),

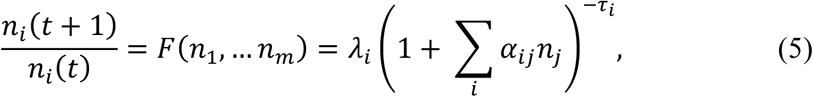

where *λ*_*i*_, > 0 is the maximum per capita seed production, *α*_*ij*_ is the per capita effect of species *j* on species *i* and *τ*_*i*_ > 0 allows each focal species to respond differently to the sum of competitor effects. This function has the partial derivative 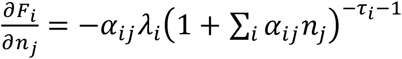. Thus, the effect of competitor *j* on the focal species *i* is a function of the density of all other competitor species. However, as in the LV model, there are no hard HOIs in this model by our definition because all of the parameters in the multi-competitor model are also found in the *m* separate single competitor functions, i.e. Θ = Φ = {*α*_*i*1_, … *α*_*im*_, *λ*_*i*_, *τ*_*i*_}.

### Why distinguish hard HOIs and non-linear density dependence (soft HOIs)?

Hard HOIs and soft HOIs have different interpretations and these differences are important to recognize if we are to advance our understanding of competition in multispecies communities. The question of whether population growth rate declines with competitor density, and whether this decline is linear or non-linear is a longstanding issue in ecology (Hassell and Comins 1976). It would be confusing at best to define HOIs as any non-linear decrease in performance with density—essentially renaming the issue of non-linear density dependence.

More importantly, hard HOIs and non-linear density dependence are ecologically distinct as well. Hard HOIs indicate a qualitative change in the way competitors affect a focal species when other competitor species are present. Non-linear density dependence, soft HOIs, does not have the same interpretation. For instance, the net outcome of competition over discrete time intervals may be non-linear when the interaction between competitors is linear in continuous time—the discrete time Hassel model, which is non-linear, is derived from a LV competition model, which is linear in continuous time (Hassell and Comins 1976, O’Dwyer 2018). In the case of the discrete time model, the lifetime competitive effect of each individual declines with competitor density because each individual competitor is smaller and thus has less of an effect on the focal species. Thus, the non-linearity in the model arguably reflects a quantitative not a qualitative change in the nature of competition when more than one species is present. In models with hard HOIs, the qualitative, or functional change in the nature of competition is defined mathematically by the introduction of additional parameters in Θ that are not present in Φ as defined above.

Adler and Morris (1994) provide another specific example where it is ecologically meaningful to differentiate between HOIs and non-linear density dependence. They describe a hypothetical scenario in which different species of plants compete for light and each species simply blocks a proportion of the light that passes through its canopy—thus taller species reduce the amount of light received by shorter species. In this way, the *qualitative* nature of the interaction between a tall species and a shorter one is independent of all other species. Nevertheless, this mechanism of interaction means that the effect of a taller species on a shorter species below it depends non-additively on the density of other competitors with a canopy between the two. Per capita competition is non-additive, but arguably there is no ecologically distinct interaction modification between the different competitors—they simply reduce the fraction of light received regardless of the presence of other species. By contrast, hard HOIs as we define them introduce new parameters, or new functional dependencies, between competitors that only kick in when more than one competitor is present.

Our definition also helps resolve the question of whether single species effects can involve HOIs. For instance, recent papers by Letten and Stouffer (2019) and Mayfield and Letten (2017) define HOIs as any higher order terms of competitor density, including single species quadratic terms, 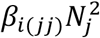. Our definition, does not count these as HOIs, and this agrees with the emphasis in the literature that HOIs are a phenomenon that arises between two or more *different species* of competitor (Hairston et al. 1968, Vandermeer 1969, Neill 1974, Morin et al. 1988). As per our definition, the coefficients for these terms, *β*_*i*(*jj*)_, are parameters in a pairwise model, *f*_*ij*_ (*n*_*j*_), and thus are not hard HOIs. Nor can single species higher order *terms* (not to be confused with higher order *interactions*) generally be interpreted as examples of *intraspecific* interaction modification, i.e. the effect of each additional individual being modified by other individuals of the same species (Mayfield and Stouffer 2017). This interpretation only makes sense in the context of a model where density dependence is strictly linear. In non-linear models, such as those fit in Mayfield and Stouffer (2017), higher order terms added to the model cannot be interpreted as individual-level interaction modifications; rather these additional terms simply allow an already non-linear function to more closely approximate the observed relationship between density and performance.

Another definition for HOIs that is largely equivalent to ours is provided by Adler and Morris (1994). Like our definition, Adler and Morris distinguished between HOIs and non-linear density dependence, and their definition agrees with ours in most cases. However, there are some cases with three or more competitor species where the Adler and Morris approach would indicate an HOI and our definition would not. We believe our definition is more general, it does not depend on the number of competitor species present and it can be more directly related to the traditional verbal definitions that ecologists have used when discussing HOIs.

In the remainder of this paper we outline the experimental set-up and statistical analyses required to test for HOIs in empirical data. Because real world data that would allow for rigorous tests of HOIs are limited, we use a mechanistic growth model to simulate a virtual competition experiment among three annual plant species (Figure 3). We then fit species’ responses to interspecific competition using phenomenological competition models with and without HOIs and evaluate which species’ responses are best fit by competition model with HOIs. By considering when HOIs emerge in this simple simulation we show the steps required to detect HOIs in empirical data and shed light on the processes that could generate HOIs in nature.

**Figure 3.**
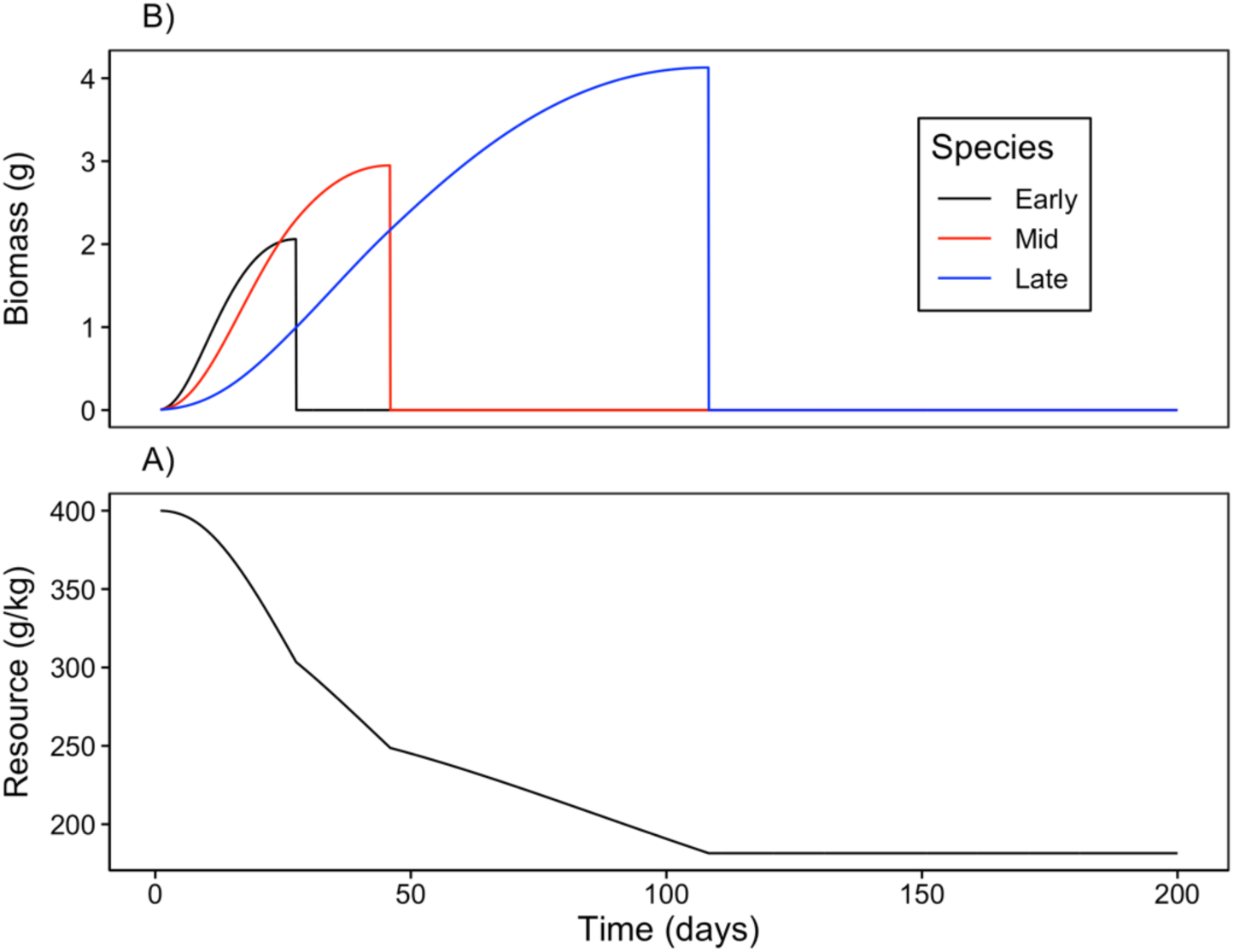
Example time course of A) annual plant growth and B) resource concentration during a single simulated growing season. In this example, each species’ population consists of a single individual. The early season species (black) grows rapidly when resource availability is high and senesces early. By contrast, the late season species (blue) grows more slowly but grows later into the season as resource availability declines. The growth curve for the mid-season species (red) lies between these extremes.

### Simulating a Higher Order Competition Experiment

A rigorous demonstration of HOIs requires measuring how focal species’ performance changes in response to increasing densities of each competitor species in isolation, as well as to varying densities of combinations of different competitor species. This requires an orthogonal response surface design where each competitor’s density is varied independently of each other species.

Instead of analyzing real data, we used a mechanistic growth model to simulate a virtual experiment in which individuals of each annual plant species are grown in separate plots with a range of competitor densities (Figure 2). The simulation lasts one growing season (200 days). After the simulation ends, we find the per capita seed output of each focal individual and record this as a measure of performance. We quantified performance in plots with densities of 0, 1, 2, 3, 4, 9, 16, 25 or 36 individuals of each other competitor species, including intraspecific competition. We also measured performance when the focal species was grown against all possible combinations of two competitor species at the same densities. This design allows us to fit non-linear functions to the interaction between each pair of species and test for any HOIs when more than two competitors are present together.

We developed a mechanistic growth and resource competition model intended to simulate the growth of annual plants in a Mediterranean climate (Figure S 1). The simulated individuals germinate in the winter and then grow, flower, and produce seeds by the early summer (Godoy and Levine 2014). In our model, we track a single pool of soil resources, most easily thought of as water or water-soluble nutrients. This pool is not resupplied during the season and is depleted over time. As the resource concentration declines, plant growth slows and eventually stops (Figure 3). We make the assumption that when individual net growth is zero, the plant will convert a fraction of its biomass into seeds that remain dormant until the start of the next growing season (Cohen 1976). Assuming all seeds germinate at the same time, and no seed mortality, we can use the per capita seed production as a direct measure of population growth rate in each competition treatment.

Resource dynamics in the model are given by the differential equation,

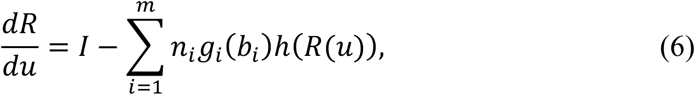

where *R*(*u*) is the resource availability at time *u* (*u* being day within the growing season), *I* is the resource supply rate, and the final term is the sum of resource uptake rates of all *m* species in the community. Biomass per individual of each species *i* at time *u* is given by *b*_*i*_ and the number of individuals in the population is given by *n*_*i*_. The function *g*_*i*_(*b*_*i*_) converts per capita biomass into surface area of fine roots. Total resource uptake rate is the product of root surface area and the rate of resource conductance per unit root surface area. The rate of resource conductance into the roots is a function, *h*(*R*), of soil resource concentration, which we specify below. We simulate a Mediterranean climate by setting initial resource availability high, *R*(*u* = 0) ≫ 0, and setting the resource supply rate, *I*, to zero.

Growth of each species is given by a piecewise differential equation,

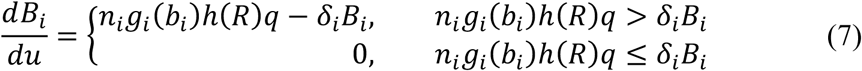

where, *q* is the rate of resource conversion into biomass and *δ*_*i*_ is the rate of biomass loss and respiration. The conditions indicate that when net growth of each species is less than or equal to zero, growth and resource consumption stops (i.e. is set to zero). Biomass per individual plant, *b*_*i*_, is converted into root surface area for each individual via the function 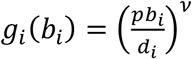, where *p* is the proportion of growth allocated to roots, *d*_*i*_ is root tissue density in g cm^−3^ and *ν* is an exponent that scales root volume to root surface area (see Kooijmans (1986) for a conceptually similar approach to protists). The rate of resource uptake per unit root surface area is dependent on resource concentration following Michaelis-Menton kinetics:

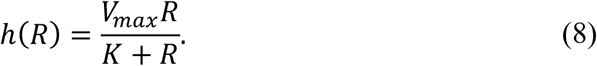

The equations above describe growth in total population biomass, *B*_*i*_, over the course of days within a single growing season. In contrast, a population-level phenomenological competition model would track the total population density, *n*_*i*_, over annual time steps, *t*. In order to convert population density into biomass, we assume that individuals start the growing season as seeds with a fixed size. Thus, the initial biomass is *B*_*i*_(0) = *μn*_*i*_(*t*), where *μ* is mass per seed and *n*_*i*_(*t*) is the number of seeds in the population in year *t*. The population density in the following year *n*_*i*_(*t* + 1) is the total number of seeds produced by the mature plants at the end of the growing season,

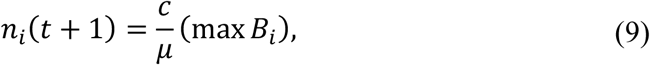

where max *B*_*i*_ is the final accumulated biomass of species *i* and *c* gives the proportion of total biomass converted to seeds.

We simulate the dynamics of three virtual annual plant species that differ in their allocation to roots and in their rates of resource uptake (Table S 1). This difference leads to phenology differences, i.e. some species stop growing earlier than others (Figure 3). Phenology differences emerge because of the assumed trade-off between species rank in terms of root density *d*_*i*_ and rank in terms of tissue respiration and loss rate, *δ*_*i*_, (Tjoelker et al. 2005, Birouste et al. 2014) (Table S1). Species with lower root density convert each gram of biomass into more root surface area and this allows them to grow faster early in the season when resource concentrations are high. In contrast, species with denser roots but lower rates of tissue loss and respiration grow more slowly but continue growing later into the season as resource availability declines. Thus, we refer to the three species in our simulations as ‘early’, ‘mid’ and ‘late’, depending on when they stop growing during the simulation (Figure 3).

We chose parameters that produced growth and phenology patterns qualitatively similar to biomass accumulation curves observed in annual plant communities (Godoy and Levine 2014). A table of parameter values for the simulations are provided in the supporting information (Table S 1). We simulated growth and resource dynamics by solving equations (6) and (7) with the package `desolve` in the statistical program R (R Core Team 2015). Code to reproduce analyses is available in a zip file and on github: https://github.com/akleinhesselink/Competitive_HOI/releases/tag/1.0

### Phenomenological annual plant model

In order to investigate whether this simulation produces HOIs between the competitors, we fit non-linear phenomenological competition models to the per capita seed production of each species. After evaluating a number of non-linear models, we found that the Hassel model (Eq. [5]) fit the outcome of simulated pairwise competition well. We specified an HOI version of the Hassel model as follows,

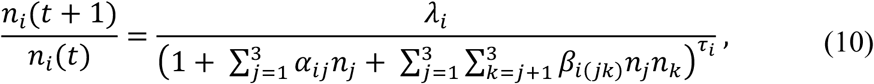

where all HOI effects of two competitor species on the focal species *i* are fitted with the coefficients *β*_*i*(*jk*)_ (following the notation in Mayfield and Stouffer (2017)). By our definition, *β*_*i*(*jk*)_ is a hard HOI when *j* ≠ *k*.

Finally, we also considered a pairwise multiplicative version of the Hassel form,

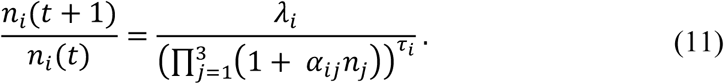

This model does not have HOIs per our definition—all *α*_*ij*_ and *τ*_*j*_ parameters can be estimated from the pairwise cases where the focal species *i* competes with each other species *j* in isolation. However, when there are two or more competitors the denominator becomes a polynomial with multiplicative terms of competitor density. In the case of only one competitor species, it collapses to the same pairwise Hassel model. Thus, contrasting this model with the HOI model allows us to test whether hard HOIs are required as opposed to a simpler non-linear function without HOIs.

We first fit the Hassel model to the pairwise cases and checked the model fit graphically. We then fit the Hassel models (Eq. [5]), the HOI model (Eq. [10]) and the multiplicative pairwise model (Eq. [11]) to the full set of two competitor densities. For each focal species and model, we calculated root mean squared error (RMSE) as a measure of goodness of fit and evaluated the strength and direction of HOIs by examining the HOI coefficients, *β*_*i*(*jk*)_. We fit all models with the non-linear least squares modelling function, `nls,` in R. Code to run the simulations, fit the models and produce the figures is given in the online supporting information.

## Results

For all three species we found the Hassel model fit the simulated pairwise data accurately (Figure 4). Next, we compared the three models fit to the full range of competitor densities (Figure 5). For the early season species, the Hassel model with and without the HOI showed more or less equivalent fits to the data with only a slight decrease in RMSE for the HOI model (Figure 5A&G). For the mid-season and late-season species, we found that the HOI model fit the data better than the pairwise Hassel model (Figure 5 B-I). The inability of the pairwise Hassel model to fit the per capita seed output of the mid and late-season species can be seen by plotting the observed and per capita seed production against two competitor densities at once (Figure S 2). In all cases, the fitted HOI coefficients, *β*_*i*(*jk*)_, were of smaller magnitude than the fitted pairwise effects, *α*_*ij*_ (Figure 6). The fitted HOIs were stronger for the mid and late season species than for the early season species (Figure 6). The multiplicative model (Eq. [11]) fit the multi-competitor dynamics poorly when compared to the pairwise model and the HOI model (Figure 5).

**Figure 4.**
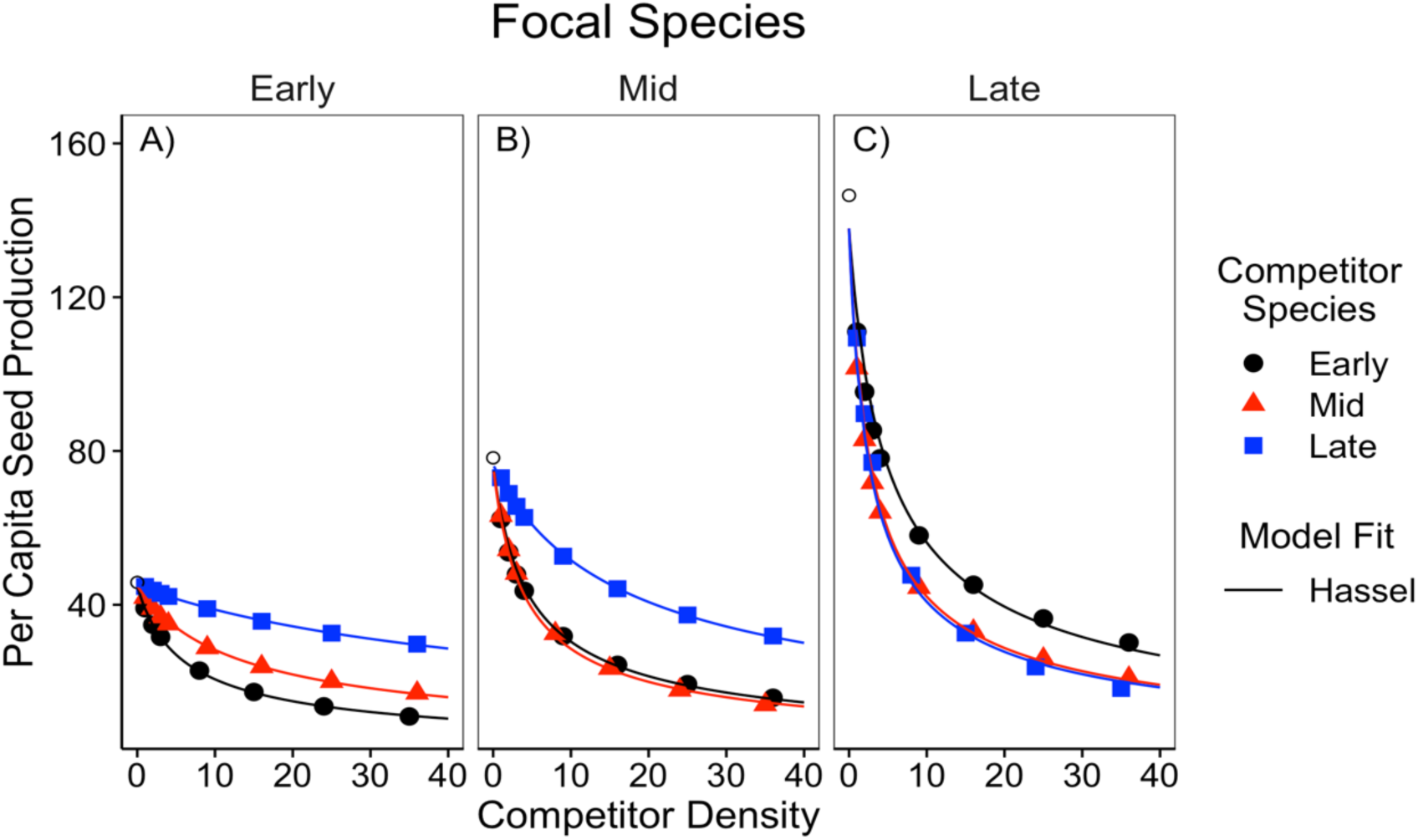
Simulated per capita seed production of the A) early, B) middle and C) late season species in response to a single competitor species at a time. Competitor density is shown on the x-axis. Colors and shapes indicate the identity of the competitor species. Open circles show the per capita seed production of each focal species in the absence of any competitors. The solid line shows the fit of the Hassel model.

**Figure 5.**
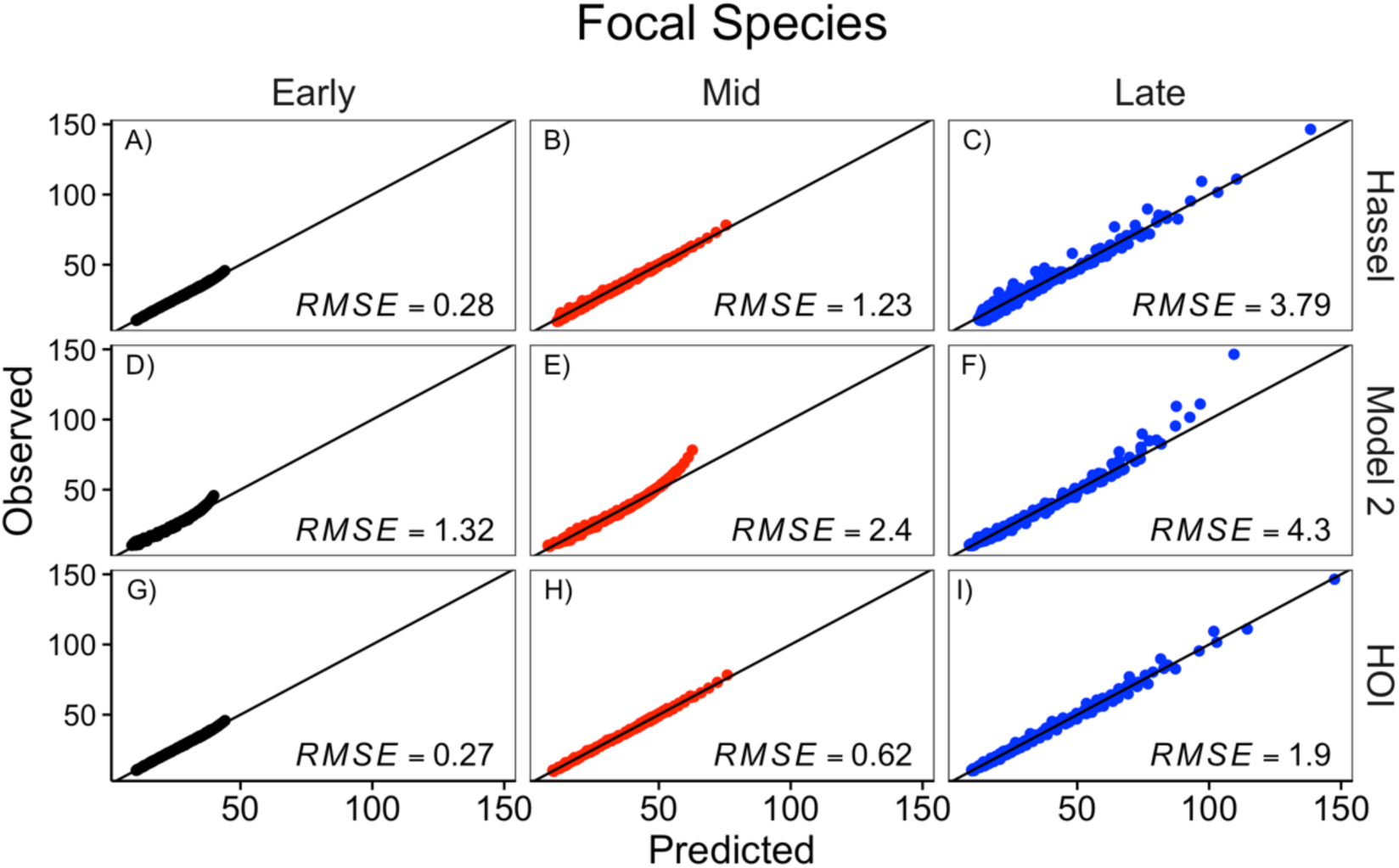
Comparison of the Hassel, multiplicative (‘model 2’), and HOI models fit to each focal species. The y-axis shows the simulated per capita seed production of the focal species. The x-axis shows the per capita seed production predicted by the phenomenological model. The top row, A-C, shows the prediction for the pairwise Hassel model (eq. [5]); the middle row, D-F, shows the prediction from the multiplicative model (eq. [11]); and the bottom row, G-I, shows the prediction from the HOI model (eq. [10]). The one-to-one line and root-mean-squared error (RMSE) for predictions from each model are shown.

**Figure 6.**
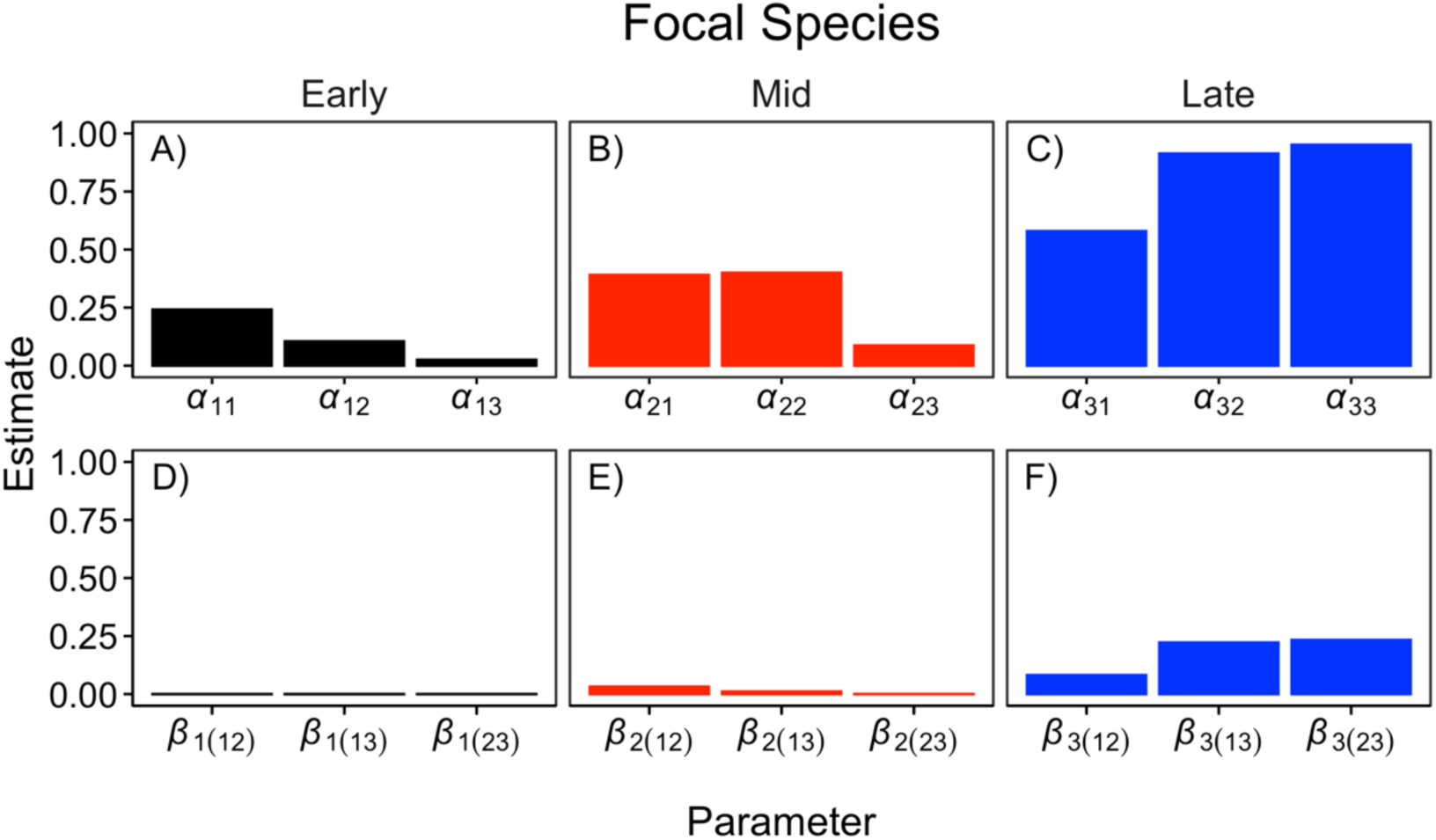
Interaction coefficients for each of focal species from the HOI model. The top row, A-C, shows the pairwise competition coefficients for the focal species and each competitor. The bottom row, D-F, shows the two-species HOI coefficients. Coefficient subscripts indicate which focal species and competitor species are involved, 1 = Early, 2 = Mid, 3 = Late.

## Discussion

### Evidence for higher order interactions

Our simulation shows clear evidence for HOIs affecting two of the three virtual species in our simulations (Figure 6). For the mid and late season species, the functional form of per capita competition changed depending on the presence of other interspecific competitors. Specifically, the presence of early or mid-season competitors increased the per capita effects of competition on the late-season species (Figure 6F). Likewise, the presence of the early season species increased the per capita effects of competition on the mid-season species (Figure 6E). For the early season species, no clear HOIs were detected: the pairwise interaction Hassel model fit the data nearly as well as the HOI model (Figure 5A&G) and the magnitudes of the HOI coefficients were small (Figure 6D).

We observe competition in our simulations because of a feedback between individual density and resource concentration. As individuals grow, they draw down resource concentrations (Figure S 1), this decreases the rate of resource acquisition into the roots by equation (8) and slows the growth of other individuals in the community. The magnitudes of pairwise interactions between species is easily understood from this perspective. For instance, the late season species has a weak per capita effect on the early season species because while the early species is active, roughly day 0 to day 30, the late-season species remains small and has a slow absolute rate of resource uptake (Figure 3A—blue line). In contrast, the mid-season species has a stronger effect on the early season species because it grows faster during the same period (Figure 3—red line). On the other hand, the early season species has a weak effect on the late season species because the former stops growing before the latter does the majority of its growth (Figure 3—black line).

The simplicity of the simulation makes it possible to understand how HOIs emerge as well. The HOIs that affect the mid and late season species are in part due to an indirect effect of resource uptake on competitor size and in part due to changes in competitor phenology. For instance, in a scenario with one individual of each species the early season species slows the growth of both the mid and the late-season species, this keeps them smaller later into the season and makes them both more sensitive to competition as the season progresses (Figure 3). This is reflected in the HOI coefficients that magnify competition for the mid and late-season species (Figure 6E&F). In contrast, the early season species grows fast and exerts the dominant effect on the resource while it is active, this makes it relatively insensitive to changes in the size of its interspecific competitors (Figure 6D).

While the HOIs in this system are similar to competition mediated indirect effects (Levine et al. 2017) there are two important differences between the HOIs we observed and traditional indirect effects. First, indirect effects are not emergent properties of a multi-competitor system, rather they are a predictable result of pairwise per capita competition coefficients (Kleinhesselink and Adler 2015). Second, indirect effects can generally be understood as emerging because of changes in the density of competitors over time, not because of changes in per capita competition. For example, one species may have an indirect effect on its competitor by changing the density of a second competitor over the course of several year. In contrast, the HOIs in our simulation emerge over the course of a single growing season with fixed population densities. Thus, these HOIs indicate ecologically meaningful changes in the per capita effect of one species on another.

Our example can be contrasted with a recent simulation of forest dynamics that demonstrated how HOIs could affect species coexistence (Grilli et al. 2017). In that simulation, unlike ours, per capita interactions between species were fixed. What the authors called HOIs in that model, were not due to changes in the per capita effect of competition, but were caused by changes in competitor density over time that were not explicitly tracked by the model.

### The phenomenological nature of HOIs

HOIs can only be defined and quantified within the context of phenomenological models of competition. Phenomenological models simplify community dynamics by tracking population densities and not the resources for which species compete (Chesson 2000). HOIs emerge in phenomenological models precisely because they leave out mechanistic detail and do not explicitly model resource dynamics (Abrams 1983, O’Dwyer 2018, Letten and Stouffer 2019). Given this, one may be tempted to conclude that HOIs are an artifact of the inadequacy of such models. However, any concept of species *interactions* (at least competitive interactions) is essentially phenomenological in nature—biomass and nutrients do not flow directly between competing individuals, rather competitors influence each other’s growth or survival indirectly through changes in the abundance of shared resources. Thus one could sidestep the problem of HOIs by instead modeling communities mechanistically as biomass and resources (e.g. Dybzinski and Tilman (2007)). However, doing may require re-thinking ecological theory formulated on the concept of species *interactions*.

Phenomenological competition coefficients can sometimes be derived analytically from mechanistic competition models by making the assumption that resource concentrations are near a fixed equilibrium (Tilman 1977, Meszéna et al. 2006, Kleinhesselink and Adler 2015, Letten et al. 2017). However, in many natural systems, such as such as those involving annual plants, resource concentrations and individual size fluctuate rapidly over the course of a single growing season or generation. This makes deriving competition coefficients directly from the resource dynamics more difficult, perhaps impossible (O’Dwyer 2018). Thus, even in cases in which we actually know which resources species compete for, fitting a phenomenological model to population dynamics may be the only way to quantitively describe species interactions. Our work clarifies the what it means to fit models with and without HOIs to multi-competitor settings.

### Are HOIs widespread?

In our virtual experiment, HOIs arise because individual size and phenology, the traits that determine each species’ impact on and sensitivity to resource availability, are themselves governed by resource availability. More generally, changes in individual size and corresponding changes in resource uptake rate may be a common cause of HOIs in nature. We predict that HOIs will likely be common in systems in which 1) consumers cause large resource fluctuations, 2) the per capita rate of resource uptake changes in response to resource availability, and 3) the strength of this response varies across species. Instead of changes in individual size, another mechanism that could generate HOIs would be density-dependent changes in resource acquisition traits. For example, traits such as height, specific leaf area, and phenology, have been shown to change in response to competition or resource availability (e.g. Aronson et al. 1992, Bennett et al. 2016, Conti et al. 2018). If per capita competition coefficients are a function of these traits, then it would not be surprising if changes in these traits led to HOIs. If changes in individual size within a season, or trait plasticity are common, and are also likely to cause HOIs, this begs the question of why there have been so few documented examples of HOIs in natural communities (but see Mayfield and Stouffer 2017).

One hypothesis is that HOIs are common but usually too weak to detect. A key factor in producing HOIs in our simulation is that each species has a uniquely shaped growth curve and phenology. In additional simulations, we found that as species became more similar in their traits HOIs became weaker (Appendix A). In nature, such large functional differences in the way species take-up resources over time may be rare. At the same time, these simulations suggest that quantifying how functional traits change in response to competitors provides a likely path to further understanding of HOIs.

A second factor generating the HOIs in our simulation are the rapid changes in resource availability and average plant size, and consequently, species interactions, over the course of a season (Figure 3). Without these dynamics, species might have relatively constant per capita effects on one another and no HOIs would emerge. For instance, compare our system to an idealized version of resource competition for perennial plants (Dybzinski and Tilman 2007). Due to their large size perennial plants can be assumed to quickly draw resources down to a dynamic equilibrium. By contrast, seasonally forced systems such as annual plant communities in Mediterranean climates may be a good place to look for strong HOIs (Mayfield and Stouffer 2017).

## Conclusion

HOIs have profound implications for how we understand and model multispecies communities. However, before ecologists can embark on measuring HOIs in nature, they must have a shared definition for what HOIs are. We have provided a more general definition of HOIs caused by interaction modifications that will be useful as ecologists seek empirical evidence for HOIs in nature. By simulating growth and resource competition in a virtual experiment, we outline the steps required to fit pairwise and HOI models to field data. This simulation also sheds light on the environmental conditions and life-history traits that may be more likely to generate HOIs. While we believe that HOIs should be common in nature this does not mean that they will be strong enough to detect statistically. Our work suggests that environments in which resource availability and competitor size change rapidly during a single growing season may be a likely place for detectable HOIs to emerge.

## Acknowledgments

We wish to thank Joanna Shih for providing the botanical line drawings. Gaurav Kandlikar, Theo Gibbs, Chris Klausmeier, two anonymous reviewers and members of the Kraft and Levine lab provided valuable comments on earlier drafts of this manuscript.

## Supporting Information – Additional Tables

**Table S 1.** Table of parameter values used in the growth simulation experiment in the main text.

| Parameter | Value | Definition |
| --- | --- | --- |
| $U$ | 200 | Duration of growth simulation (days) |
| $I$ | 0 | Resource supply rate ($\text{g day}^{-1}$) |
| $R(0)$ | 400 | Initial resource concentration ($\text{g kg}^{-1}$) |
| $d_1$ | 0.06 | Early competitor root density ($\text{g cm}^{-3}$) |
| $d_2$ | 0.12 | Mid competitor root density ($\text{g cm}^{-3}$) |
| $d_3$ | 0.36 | Late competitor root density ($\text{g cm}^{-3}$) |
| $\delta_1$ | 0.3 | Early competitor loss and respiration rate ($\text{g g}^{-1}\text{d}^{-1}$) |
| $\delta_2$ | 0.15 | Mid competitor loss and respiration rate ($\text{g g}^{-1}\text{d}^{-1}$) |
| $\delta_3$ | 0.053 | Late competitor loss and respiration rate ($\text{g g}^{-1}\text{d}^{-1}$) |
| $K$ | 350 | Resource half-saturation constant ($\text{g kg}^{-1}$) |
| $V_{max}$ | 1 | Maximum resource conductance ($\text{g d}^{-1}\text{cm}^{-2}$) |
| $p$ | 0.5 | Ratio of root biomass to total biomass |
| $\nu$ | 0.66 | Scaling exponent (unitless) |
| $q$ | 0.2 | Biomass assimilation rate ($\text{g g}^{-1}\text{d}^{-1}$) |
| $\mu$ | 0.005 | Seed mass (g per seed) |
| $c$ | 0.1 | Conversion of final biomass to seed mass ($\text{g g}^{-1}$) |

## Supporting Information – Additional figures

**Figure S 1.**
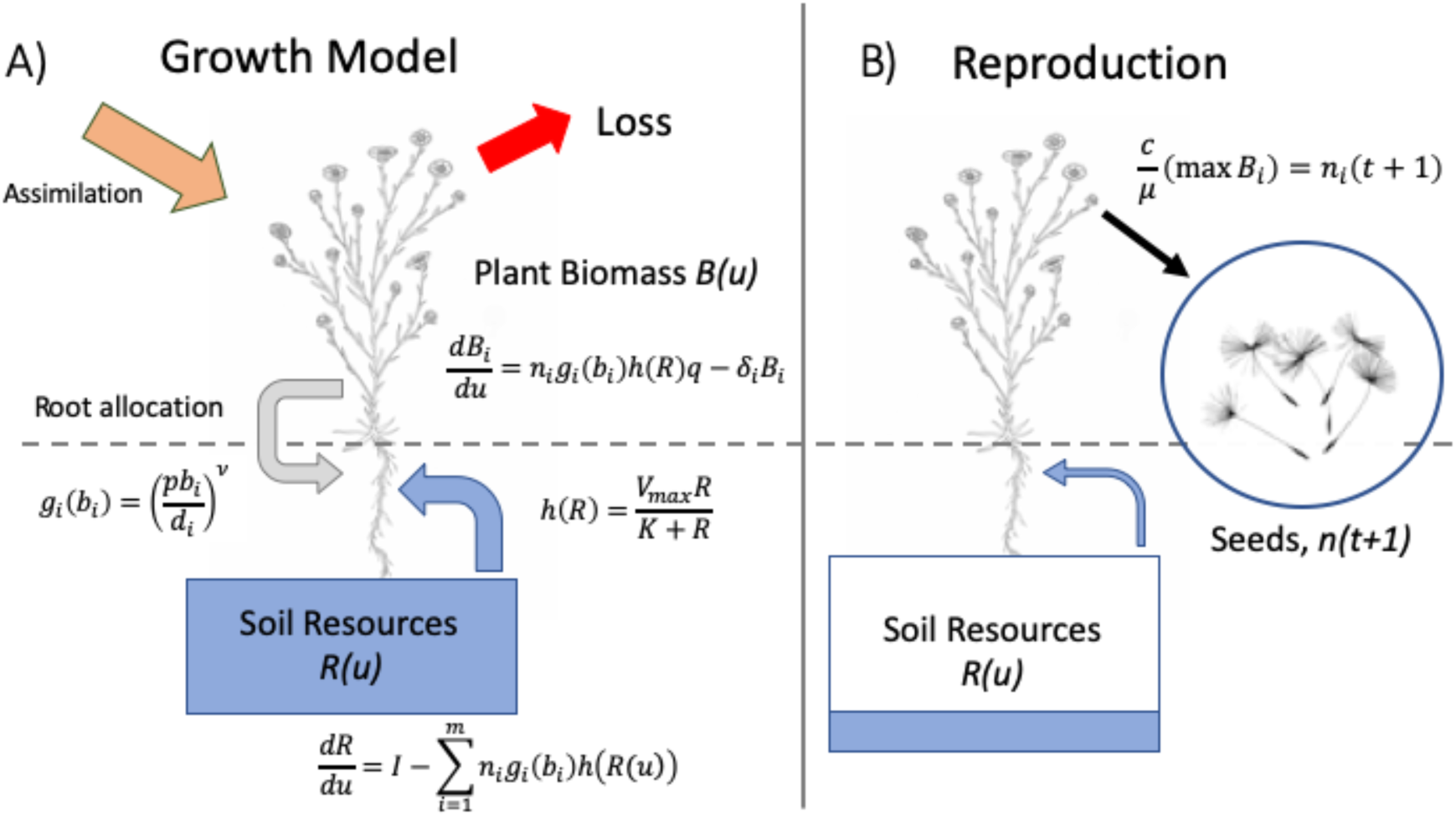
Diagram schematic of annual plant growth model used in simulation. A) in the model each individual plant start as a seed, grows over the course of a single growing season. Growth is a function of plant biomass, root surface area and soil resource availability. B) Over time the soil resources are depleted and plant growth slows down. Plants reach a maximum size when losses due to respiration and tissue senescence are greater than growth. At this point the plants convert stored resources to seeds. The number of seeds in the next growing season is determined as the total mass of seeds produced per species divided by the weight of a single seed.

## Supporting Information – Additional figures

**Figure S 2.**
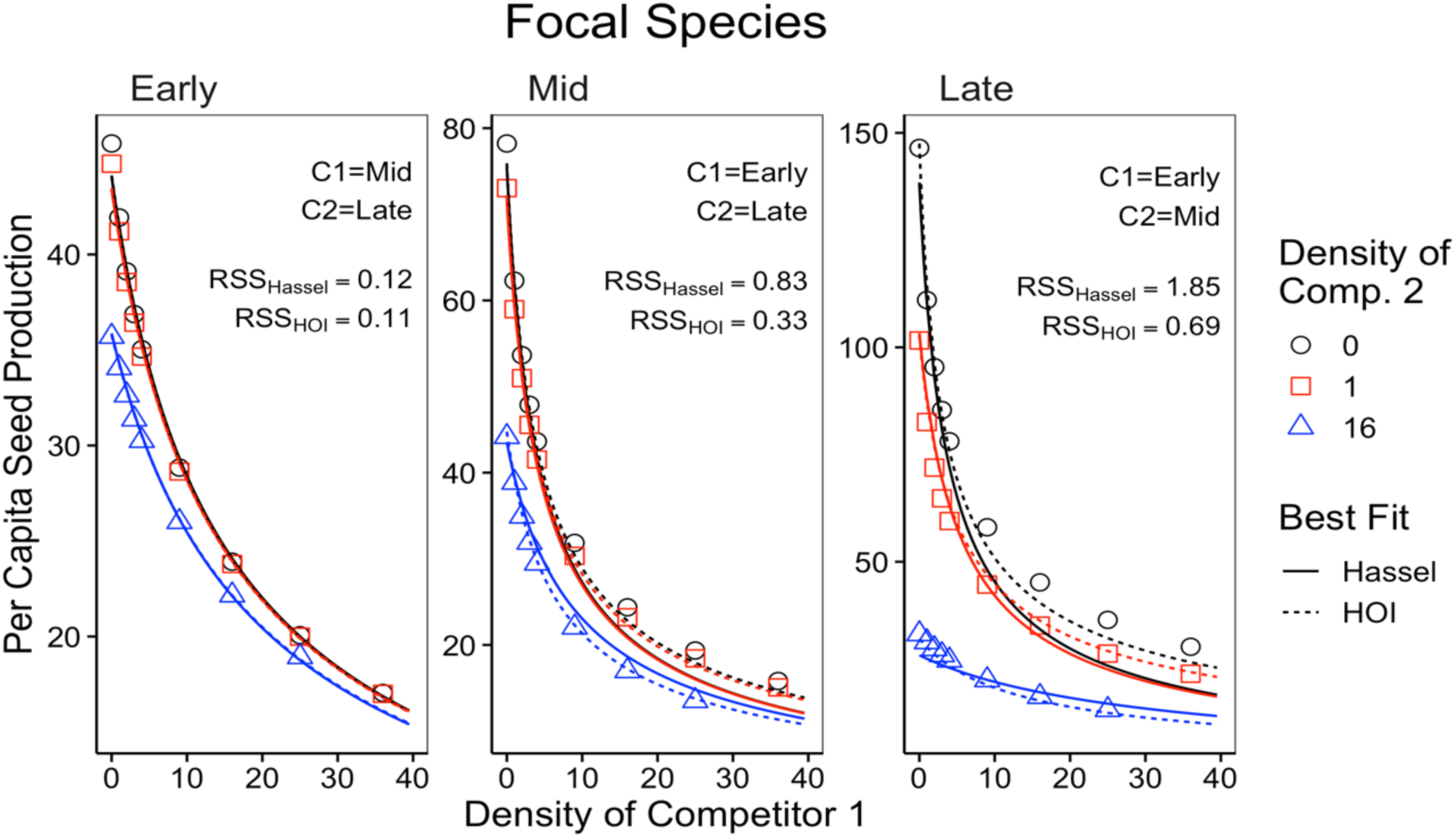
Simulated per capita seed production of the A) early, B) mid and C) late season species in response to density of two interspecific competitors. Densities of two competitors are shown in each panel—the x-axis shows the density of the first competitor, while different colored lines and shapes show the density of a second competitor. Text in each panel lists the identities of competitor one and two (early, mid or late). Lines show best fit from phenomenological models. Residual sum of squared error is shown for each model and focal species.

### Appendix A – The effect of trait differences on higher order interactions

We used an additional simulation experiment to test whether the strength of higher order interactions (HOIs) was associated with the magnitude of functional differences between competitor species. We started with the same parameter values as in the simulation in the main text in which there was a large difference between the species in root density (*d*_*i*_) and tissue respiration rate (*δ*_*i*_). In four additional simulation scenarios, we gradually decreased the average difference between species in these traits (Table A1). Specifically, we held the traits of the mid-season species constant and decreased the difference in the root density trait, *d*_*i*_, between the early and late-season species. We assumed a trade-off between root density and tissue respiration rate such that changing root density was accompanied by a change in tissue respiration rate, *δ*_*i*_ (Figure A1). We quantified the average functional difference between species as the standard deviation of root density among all species. In each scenario, we simulated competition and fitted the phenomenological HOI model as in the main text. For each species in each scenario, we quantified the strength of HOIs as the average magnitude of the *β* coefficients divided by the average magnitude of the *α* coefficients. For the mid and late season species, the strength of the HOIs increased with the functional difference between species (Figure A1 B&C). For the early season species, HOIs were weak in all five scenarios (Figure A1 A). These simulations show that the functional differences between competitors drive the HOIs we observed in this system.

**Table A 1.**
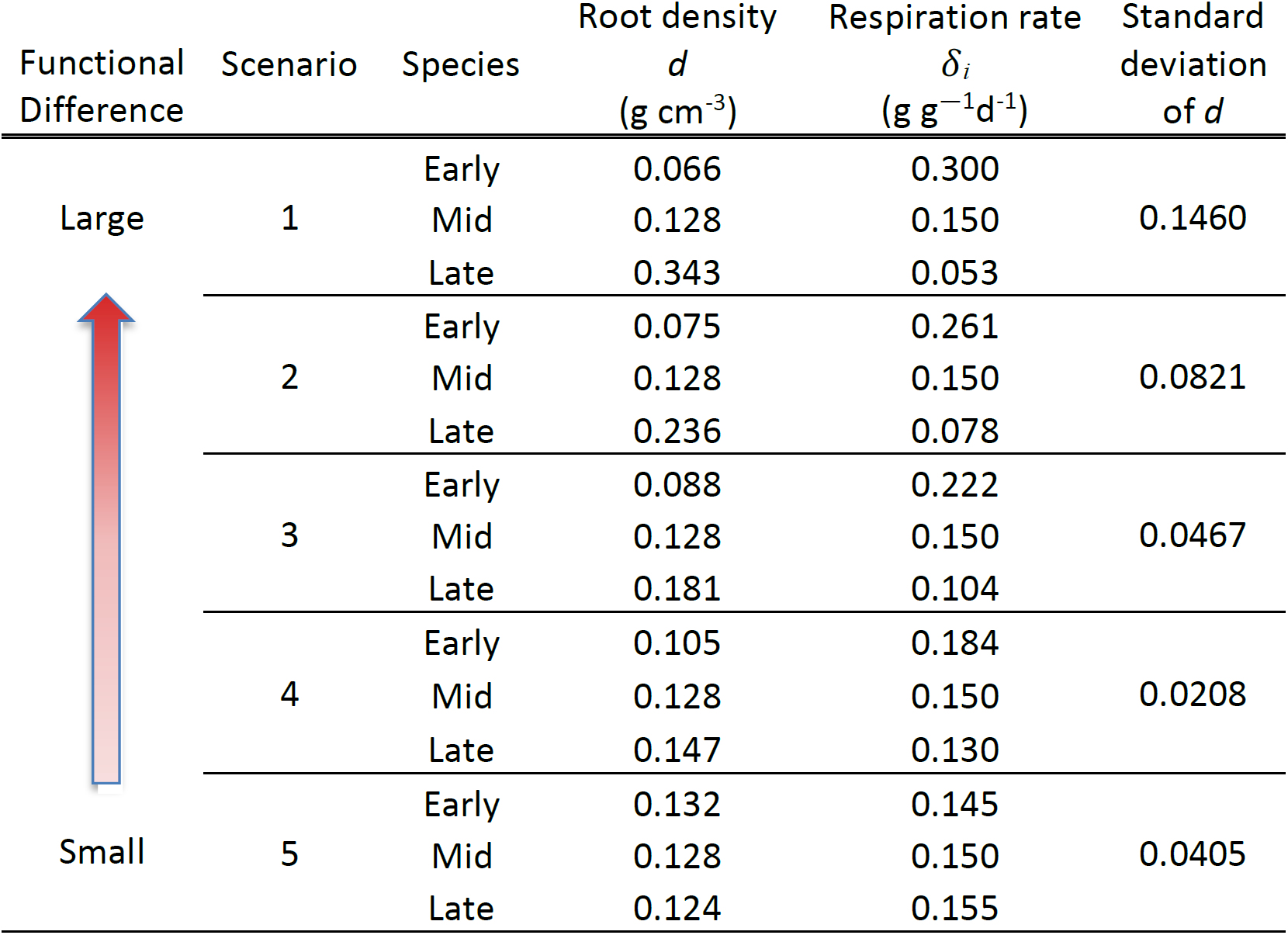
Parameter values for five simulations with gradually decreasing the trait difference between the early season and late season species. All other simulation parameters are the same as in Table S1.

**Figure A 1.**
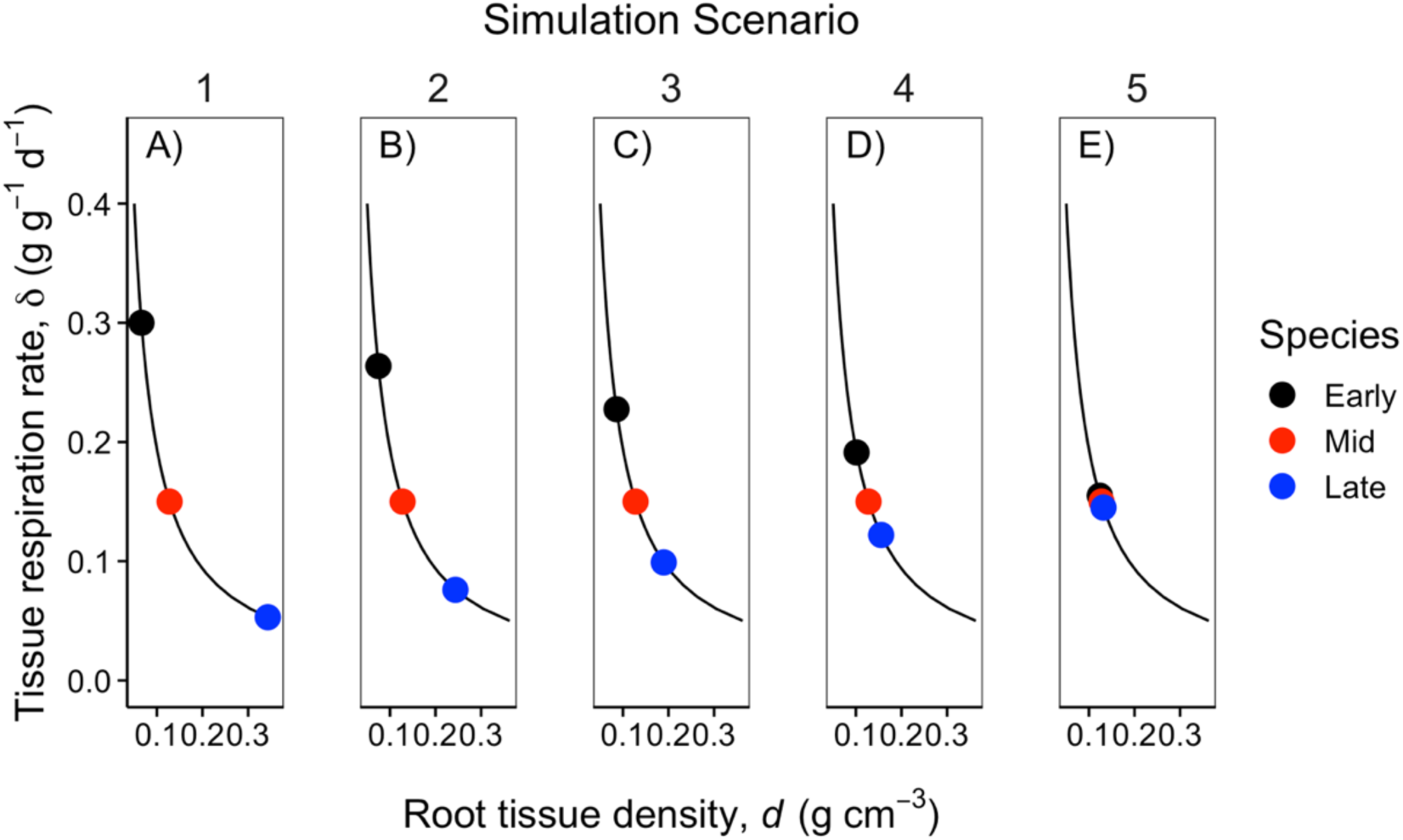
Colored points show the value of functional traits, root density and tissue loss rate, for each of the three species in each of the five simulation scenarios (A-E). Across the five scenarios, the differences between the early season and late season species’ root density and respiration rates were gradually decreased. The mid-season species’ traits were held constant. The black line indicates the trade-off between the root density and tissue respiration rate traits.

**Figure A 2.**
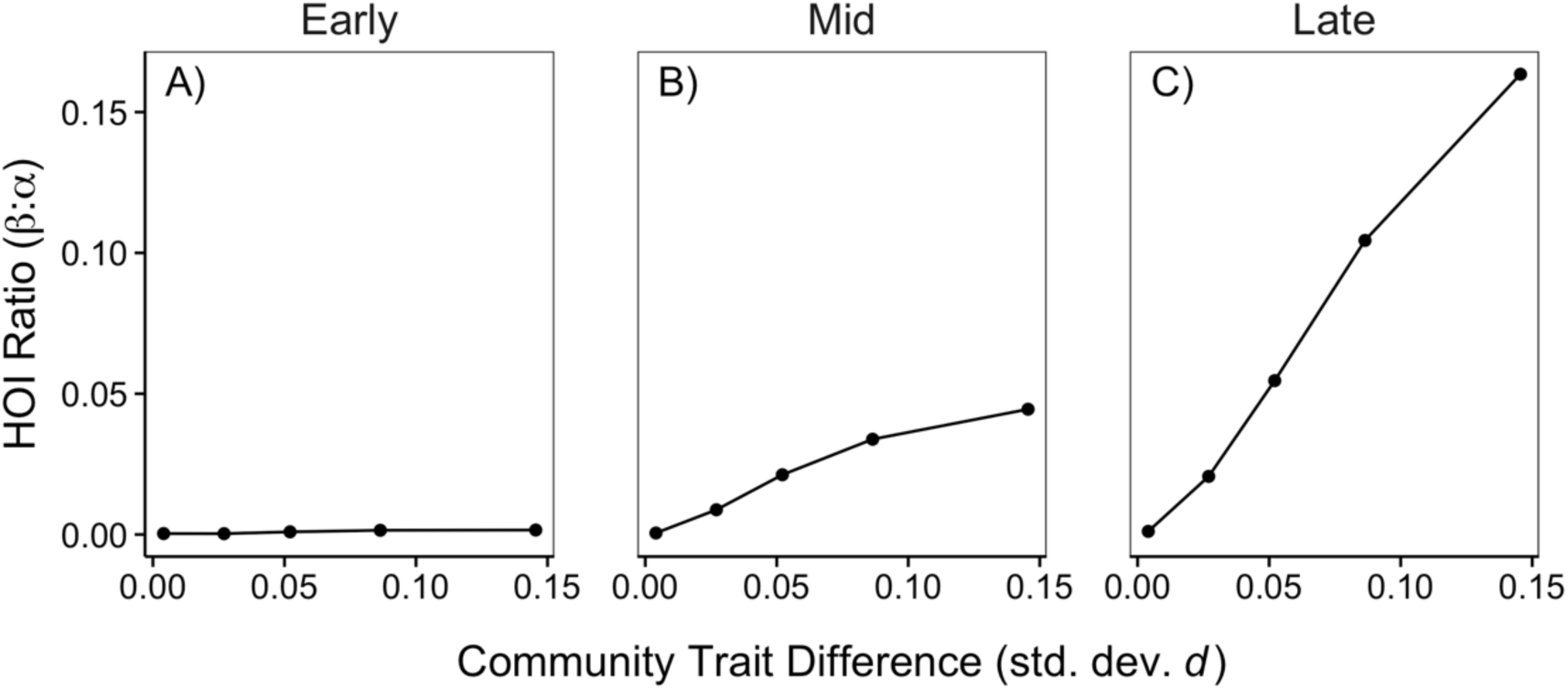
The strength of HOIs depends on the difference in species functional traits. The y-axis quantifies the strength of HOIs affecting the early (A), mid (B) and late (C) species as the ratio of the of the average magnitude of the *β*_*i*(*jk*)_ coefficients to the average magnitude of the *α* coefficients in the phenomenological HOI model. A larger ratio *β*: *α* ratio indicates stronger HOIs compared to pairwise interactions. The x-axis quantifies the community-level trait difference as the standard deviation of the root density trait, *d*.

